# Evaluating Sleep Disturbances in Children with Rare Genetic Neurodevelopmental Syndromes

**DOI:** 10.1101/2021.02.12.430633

**Authors:** Olivia J. Veatch, Beth A. Malow, Hye-Seung Lee, Aryn Knight, Judy O. Barrish, Jeffrey L. Neul, Jane B. Lane, Steven A. Skinner, Walter E. Kaufmann, Jennifer L. Miller, Daniel J. Driscoll, Lynne M. Bird, Merlin G. Butler, Elisabeth M. Dykens, June-Anne Gold, Virginia Kimonis, Carlos A. Bacino, Wen-Hann Tan, Sanjeev V. Kothare, Sarika U. Peters, Alan K. Percy, Daniel G. Glaze

**Affiliations:** Department of Psychiatry and Behavioral Sciences, University of Kansas Medical Center, Kansas City, KS, USA; Vanderbilt Kennedy Center, Departments of Pediatrics and Neurology, Vanderbilt University Medical Center, Nashville, TN, USA; Department of Pediatrics University of South Florida, Tampa, FL, USA; Center for Clinical Research, Texas Heart Institute, Houston, TX, USA; Departments of Pediatrics and Neurology, Baylor College of Medicine, Houston, TX, USA; Vanderbilt Kennedy Center, Departments of Pediatrics, Pharmacology, and Special Education, Vanderbilt University Medical Center, Nashville, TN, USA; University of Alabama at Birmingham, School of Medicine, Birmingham, AL, USA; University of Alabama at Birmingham, Civitan International Research Center, Birmingham, AL, USA; Center for Translational Research, Greenwood Genetic Center, Greenwood, SC, USA; Department of Neurology, Boston Children’s Hospital, Boston, MA, USA; Department of Pediatrics, University of Florida, Gainesville, FL, USA; Department of Pediatrics, Division of Genetics and Dysmorphology, University of California San Diego/Rady Children’s Hospital, San Diego, CA, USA; Vanderbilt Kennedy Center, Departments of Pediatrics and Special Education, Vanderbilt University Medical Center, Nashville, TN, USA; Division of Genetics and Genomic Medicine, Department of Pediatrics, University of California, Irvine, CA, USA; Division of Genetics and Genomics, Boston Children’s Hospital, Boston, MA; Pediatric Sleep Program, Cohen Children’s Medical Center, New Hyde Park, New York; Vanderbilt Kennedy Center, Departments of Pediatrics and Psychiatry & Behavioral Sciences, Vanderbilt University Medical Center, Nashville, TN, USA

**Author notes:** Corresponding Author: Olivia J. Veatch, PhD, Department of Psychiatry and Behavioral Sciences, University of Kansas Medical Center, 3901 Rainbow Blvd, Kansas City, KS, 66160. Current Affiliations: Department of Human Genetics, Emory University School of Medicine, Atlanta, GA, USA and Anavex Life Sciences Corp., New York, NY, USA.

## Abstract

**Background:** Adequate sleep is important for proper neurodevelopment and positive health outcomes. Sleep disturbances are more prevalent in children with genetically determined neurodevelopmental syndromes compared to typically developing counterparts. We characterize sleep behavior in Rett (RTT), Angelman (AS) and Prader-Willi (PWS) syndromes in order to identify effective approaches for treating sleep problems in these populations. We compared sleep-related symptoms across individuals with these different syndromes to each other, and to typically developing controls.

**Methods:** Children were recruited from the Rare Diseases Clinical Research Network (RDCRN) consortium registries; unaffected siblings were enrolled as related controls. For each participant, a parent completed multiple sleep questionnaires including: Pediatric Sleep Questionnaire (Sleep-Disordered Breathing [SDB]); Children’s Sleep Habits Questionnaire; Pediatric Daytime Sleepiness Scale.

**Results:** Sleep data were analyzed from 714 participants, ages 2-18 years. Young children with AS had more reported sleep problems than children with RTT or PWS. Older children with RTT had more reported daytime sleepiness than those with AS or PWS. Finally, all individuals with RTT had more evidence of sleep-disordered breathing when compared to individuals with PWS. Notably, typically developing siblings were also reported to have sleep problems, except for sleep-related breathing disturbances which were associated with each of the genetic syndromes.

**Conclusions:** Individuals with RTT, AS and PWS frequently experience sleep problems, including sleep-disordered breathing. Screening for sleep problems in individuals with these and other neurogenetic disorders should be included in clinical assessment and managements. These data may also be useful in developing treatment strategies and in clinical trials.

## Introduction

Quality and quantity of sleep is important for optimal human health, as evidence indicates sleep influences many physiological and behavioral functions^1^. Critical roles for sleep in learning, memory, neurogenesis, and neural plasticity have been described, emphasizing the importance of sleep to brain development and function^2–6^. This suggests that healthy sleep in children is essential for appropriate neurodevelopment and ultimate neurological function. The range of sleep disorders in children is broad, although the details of how different sleep problems impact typical development and chronic illness remain unclear. Sleep disturbances have severe adverse effects on the quality of life of children and their families^7^. In the general pediatric population, sleep problems affect approximately 25% of all children^2^. In neurodevelopmental disorders, 50-95% of individuals may experience sleep problems that constitute a risk for exacerbation of daytime problematic behaviors and poor cognitive and academic performance^4,7–9^. Of the more than 1,200 individuals with genetically determined neurodevelopmental syndromes enrolled in the NIH funded Rare Diseases Clinical Research Network (RDCRN) Rett, Angelman and Prader-Willi syndromes consortiums, most have sleep problems. These problems include difficulty initiating and maintaining sleep, sleep-disordered breathing, and daytime sleepiness^10^. However, little information is available regarding whether a specific sleep profile is observed in distinct genetic syndromes and whether these sleep problems are persistent and chronic. The natural history studies of Rett, Angelman and Prader-Willi syndromes conducted as part of the RDCRN provide the opportunity to examine sleep profiles in a large cohort of rigorously molecularly diagnosed and comprehensively clinically characterized individuals over time.

Rett syndrome (RTT) is a neurodevelopmental disorder that primarily affects females with an incidence of 1/10,000–1/15,000^11–15^. RTT is characterized by an initial period of apparently typical development followed by a period of regression with loss of spoken communication and purposeful hand use, onset of stereotypical hand movements and impairment or absence of ambulation^16^. Following the regression phase, a period of stabilization without further cognitive decline is noted, though motor problems may worsen, and other manifestations arise. The latter include epilepsy, growth failure, gastrointestinal issues (e.g., constipation and gastrointestinal esophageal reflux), scoliosis, and autonomic dysfunction. While life expectancy is shorter than in the general population, recent data demonstrate survival to the sixth decade in most individuals with RTT^17^. The majority of cases of RTT are caused by pathogenic variants leading to loss of function in the X–linked gene methyl–CpG–binding protein 2 (*MECP2*)^11–13,16,18^. Notably, sleep problems are frequently reported in RTT individuals of all ages and are one of the supportive criteria for clinical diagnosis^16,19,20^.

Angelman syndrome (AS) is a neurogenetic disorder affecting both females and males, with a prevalence of 1/22,000 – 1/52,000^21–24^. The cause of AS is the absence of functional UBE3A protein in neurons related to a deletion of the maternal chromosome 15q11-q13 which includes the imprinted maternally expressed ubiquitin protein ligand E3A (*UBE3A*) gene. The phenotype, present from birth, is characterized by severe developmental impairment, absent or minimal spoken language, gait abnormalities, happy and excitable personality, epilepsy, and abnormal movements. Children with AS frequently experience insomnia-related symptoms and appear to sleep less than age-matched typically developing children. Sleep problems are reported to improve with age^25,26^; however, it remains unclear if this finding is consistent among all AS patient populations.

Prader-Willi syndrome (PWS) is a neurodevelopmental disorder affecting both males and females, with an incidence between 1/10,000 to 1/30,000^27–29^. The molecular cause is due to loss of the paternally imprinted region on chromosome 15q11-q13. The majority of cases occur as a result of a 15q11-q13 deletion involving the paternal chromosome 15, followed by maternal disomy of chromosome 15, and imprinting center defects^26,30^. The phenotype in infancy is characterized by hypotonia, feeding difficulties, poor weight gain, distinctive facial features, and developmental delay. In childhood patients develop hyperphagia, which in combination with low metabolic rate, often results in early-onset morbid obesity^31^. Sleep problems occur in most individuals with PWS and are characterized by obstructive or central sleep apnea (with some individuals experiencing both) and hypersomnia may occur in the absence of sleep-disordered breathing; some patients also have findings consistent with narcolepsy^32–34^.

We leveraged parent reports of sleep behaviors in individuals who participated in the RDCRN natural history studies to better characterize sleep problems in one of the largest cohorts of individuals with RTT (NCT02738281), AS (NCT00296764) and PWS (NCT03718416) to date. The goals were to identify sleep patterns that were common among children and adolescents with these syndromes and determine distinctions among these three syndromes. Given that each of these syndromes have well-described unique genetic causes, it is possible that these data may inform future work aimed at understanding genetic contributions to sleep problems in these patients. Additionally, we compared data from syndromic children and their siblings, to explore the effects of the affected childrens’ sleep problems on their typical siblings.

## Materials and Methods

### Study design

Individuals were recruited from the RDCRN natural history studies for AS, RTT, PWS. These studies collect clinical and genetic diagnosis, medical history and contact information. The RDCRN consortium sites recruited subjects, obtained informed consent and administered sleep questionnaires. Institutional Review Board approval was obtained at each site and included: University of Alabama at Birmingham (RTT), Baylor College of Medicine (RTT, AS), Boston Children’s Hospital (RTT, AS), Greenwood Genetic Center (RTT, AS), Rady Children’s Hospital San Diego (AS), Vanderbilt University (AS, PWS), University of California – Irvine (PWS), University of Florida (PWS), and University of Kansas Medical Center (PWS).

All individuals enrolled in the sleep project were required to be ≤19 years old at the time of enrollment, live in a private home with a parent or guardian familiar with the child’s specific sleep pattern, and the parent or guardian was required to be fluent in English (study questionnaires were only available in English). Individuals were excluded if they lived in a group home where the parent/guardian was not the primary caregiver. Affected children were defined as individuals with a diagnosis of RTT, AS, or PWS. Unaffected children were those who had a sibling with either RTT, AS, or PWS enrolled in the study, who did not themselves have a diagnosis of any neurological disorder. For each participant, a parent/caregiver who was familiar with the sleep habits completed each of the study questionnaires. Questionnaires were checked for completeness; parents/caregivers were requested to complete unfinished items. Multiple questionnaires were used to ensure a comprehensive assessment of sleep-related traits.

We considered including data from individuals of all ages enrolled via the RDCRN but since we incorporated comparisons of data from a typically developing sibling sample, limited our analyses to data from participants within age ranges where questionnaires have been validated. We analyzed data from the following questionnaires:

1. The **Children’s Sleep Habits Questionnaire (CSHQ)** is a retrospective, 45-item parent questionnaire that has been used and validated in a number of studies to examine sleep behavior in young children, from 2-10 years old^35,36^. To calculate a total score, answers to 33-items are summed with higher scores indicating more problematic sleep behaviors. Scores can range from 33-99 with scores of 41 or greater suggested to indicate the significant occurrence of all types of sleep problems^35^. We analyzed total scores from the CSHQ for children between the ages of 2-10 years old.
2. The **Pediatric Daytime Sleepiness Scale (PDSS)** includes eight questions to assess daytime sleepiness and is similar to the Epworth Sleepiness scale for adults^37^. The range of possible scores is 0-32. The Pediatric Daytime Sleepiness Scale questionnaire has been validated for assessment of sleep in individuals between the ages of 5-17 years^37–39^. We analyzed total scores for daytime sleepiness for participants between the ages of 5-17 years old.
3. The **Sleep-Related Breathing Disorder Scale (SRBD)** consists of 22 closed response question-items, extracted from the **Pediatric Sleep Questionnaire (PSQ)** which was developed for clinical research purposes, and validated against polysomnography^40^. Final scores for the SRBD scale are based on the mean response to all relevant questions where ‘yes’ = 1 and ‘no’=0. The PSQ is validated for individuals between the ages 2-18 years; we analyzed SRBD scores for all participants within this age range.

Although score ranges differ, it is notable that for all instruments higher total scores indicate more reported sleep problems/increased severity.

### Statistical Analysis

Age was determined by calculating the difference between the birthdate and date of the visit where the parent completed questionnaires. For a subset of individuals (n=48), the birthdate was missing or the calculated age at the visit was improbable (i.e., negative or <1 year). Missing or improbable ages were replaced with the reported age at registration when available; when age could not be determined, individuals were excluded. In addition, to comply with the validity of the questionnaires, individuals who were younger than 2 years old or older than 18 years old were excluded. To assess normality assumptions, histograms for ages and sleep questionnaires were plotted and Shapiro-Wilk’s tests conducted. Kruskal-Wallis tests were used to compare age and proportion tests to compare gender and reported ethnicity across all participants. For subjects with any of the syndromes, Kruskal-Wallis tests were used to compare total scores from the sleep questionnaires. Significant results were evaluated using post-hoc Dunn tests and p-values were adjusted with the Benjamini-Hochberg method. Conditional logistic regression was used, while accounting for relatedness, to first determine if age was associated with any scores and then to compare parent reports of sleep problems between the individual with the genetic syndrome to their sibling control(s). Additional models adjusting for age at visit were evaluated given that age was associated with the measure. Mean and standard deviation are reported. All reported p-values are two-sided with a p-value<0.05 considered statistically significant. All analyses were performed using R v 3.5.1 (The R Foundation for Statistical Computing, Vienna, Austria).

## Results

### Study Population

The overall enrollment was 792 individuals. Of the total enrollment, data from 714 participants with available ages between 2-18 years old whose parents had completed at least one sleep questionnaire were analyzed. The largest cohorts were the sibling controls (N=293), followed by individuals diagnosed with RTT (N=246), then PWS (N=95), or AS (N=80). The analysis dataset was predominantly white (87%), reflecting the ethnicity of the RDCRN participants in general.

Gender was majority female (70%) and the proportion was different across groups, reflecting the large number from the RTT cohort which was almost exclusively female (∼99%) as the *MECP2* gene is X-linked; gender was not different between patients with AS or PWS (p=0.66). Age was significantly different from a normal distribution (Figure 1; W=0.97, p-value<0.0001). The mean age ranged from 7.77 to 9.76 years with significant differences when comparing all diagnostic groups, including unaffected siblings (Table 1); however, no age differences were observed among the three genetic syndromes (p=0.56).

**TABLE 1:** Study participants with at least one sleep questionnaire completion.

| Diagnosis | Age in Years<br>Mean±SD (Range) | Ethnicity<br>% White | Total N<br>(% Female) |
| --- | --- | --- | --- |
| Rett syndrome (RTT) | 9±4 (2-17) | 83 | 246 (99) |
| Sibling controls (RTT) | 10±4 (2-18) | 86 | 224 (59) |
| Angelman syndrome (AS) | 8±4 (2-18) | 90 | 80 (53) |
| Sibling controls (AS) | 8±4 (2-18) | 90 | 29 (41) |
| Prader-Willi syndrome (PWS) | 9±4 (2-18) | 88 | 95 (57) |
| Sibling controls (PWS) | 8±4 (2-18) | 95 | 40 (48) |
| *p-value†* | 0.01 | 0.31 | <0.0001 |
| **Total** | 9±4 (2-18) | 87 | 744 (71) |

†p-values for tests of differences comparing age, ethnicity and gender among diagnostic groups.

**Figure 1.**
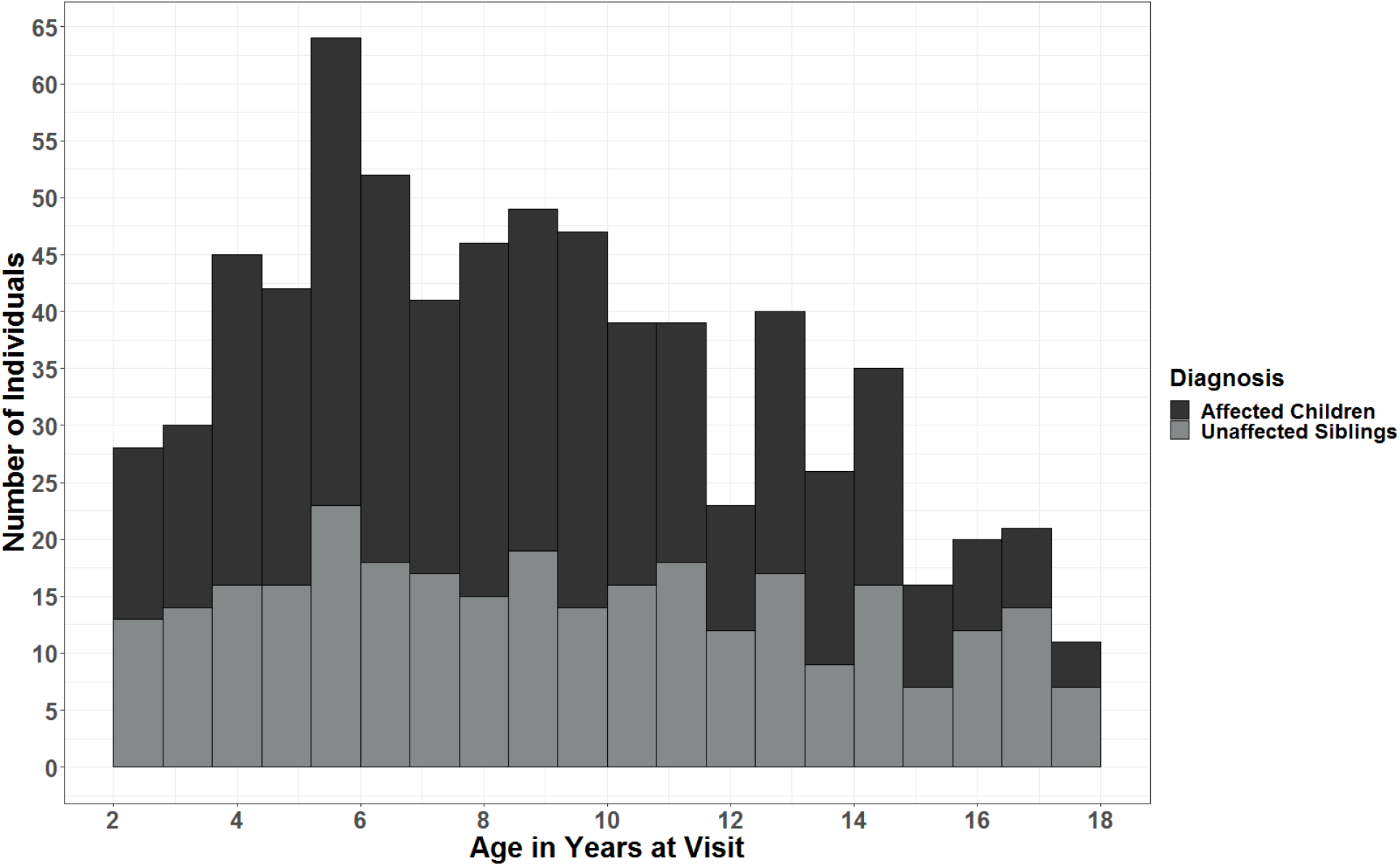
Age distribution for individuals with sleep data included in analyses. Shown are ages in years of children affected with a genetic syndrome (dark gray) and their unaffected siblings (light gray), at the time of parent sleep questionnaire completion.

### Comparison of Sleep Disturbances among Different Genetic Syndromes

Results from comparisons of parent-reported sleep for individuals with each evaluated genetic syndrome are provided in Table 2. Score distributions for all questionnaires were not normal (p-value<0.0001). All children were reported to have significant sleep problems based on CSHQ score cut-offs (>43). Specifically, children with AS had higher total CSHQ scores when compared to children with RTT or PWS and children with RTT had higher scores than those with PWS. Older children and adolescents with RTT were reported to have more issues with daytime sleepiness on the PDSS when compared to both AS and PWS. No differences were observed when comparing PDSS between individuals with AS and PWS. Finally, individuals across all ages with RTT and AS showed no differences in parent-reported symptoms of sleep-disordered breathing on the PSQ; however, individuals with RTT had higher scores when compared to individuals with PWS.

**TABLE 2A:**
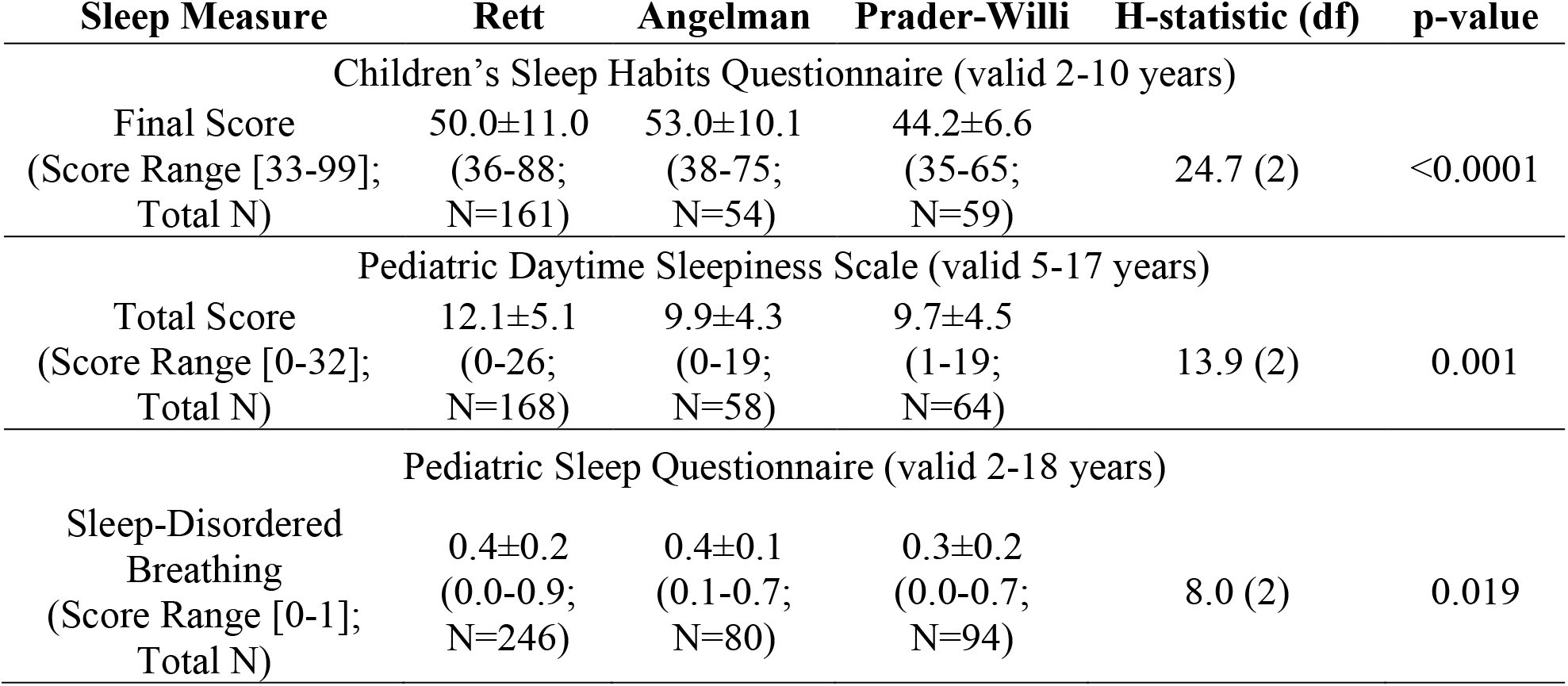
Differences in sleep problems among genetic syndromes.

**TABLE 2B:** Direct comparisons of sleep problems between genetic syndromes.

| **Sleep Measure** | **Genetic Syndromes Comparison** | **z-statistic** | **p-value<sub>adjusted</sub>** |
| --- | --- | --- | --- |
| CSHQ Total | RTT vs AS | -2.3 | 0.02 |
|  | RTT vs PWS | 3.7 | 0.0004 |
|  | AS vs PWS | 4.9 | <0.0001 |
| PDSS Total | RTT vs AS | 2.9 | 0.01 |
|  | RTT vs PWS | 3.3 | 0.003 |
|  | AS vs PWS | 0.5 | 0.63 |
| SDB Score | RTT vs AS | 0.3 | 0.78 |
|  | RTT vs PWS | 2.8 | 0.02 |
|  | AS vs PWS | 2.0 | 0.07 |

Reported are results of Kruskal-Wallis (A) and Dunn (B) tests for differences in parent-reported sleep problems among the three genetic syndromes. For each questionnaire, the valid age and possible score ranges are noted. Means±standard deviations are provided, along with the range of scores observed and total number of patients (N) whose data were analyzed. Dunn test p-values are Benjamini-Hochberg adjusted. H-statistics reflect tests of whether medians are equal, z-statistics reflect pairwise multiple comparisons based on rank sums. df=degrees of freedom.

### Comparison of Sleep Disturbances between Genetic Syndromes and Sibling Controls

Increases in parent reported sleep-related breathing disturbances on the PSQ were associated with having a diagnosis for each of the genetic syndromes evaluated when compared to siblings (Table 3). There was no evidence for associations between total scores on the CSHQ or daytime sleepiness reported on the PDSS and having a genetic syndrome. Notably, scores on the PDSS were associated with age in the families with Rett Syndrome (p=0.03). When age was included in the regression model, the association between increased scores on the PDSS and a diagnosis of Rett syndrome was nearly significant (p=0.05; Table 3).

**TABLE 3.**
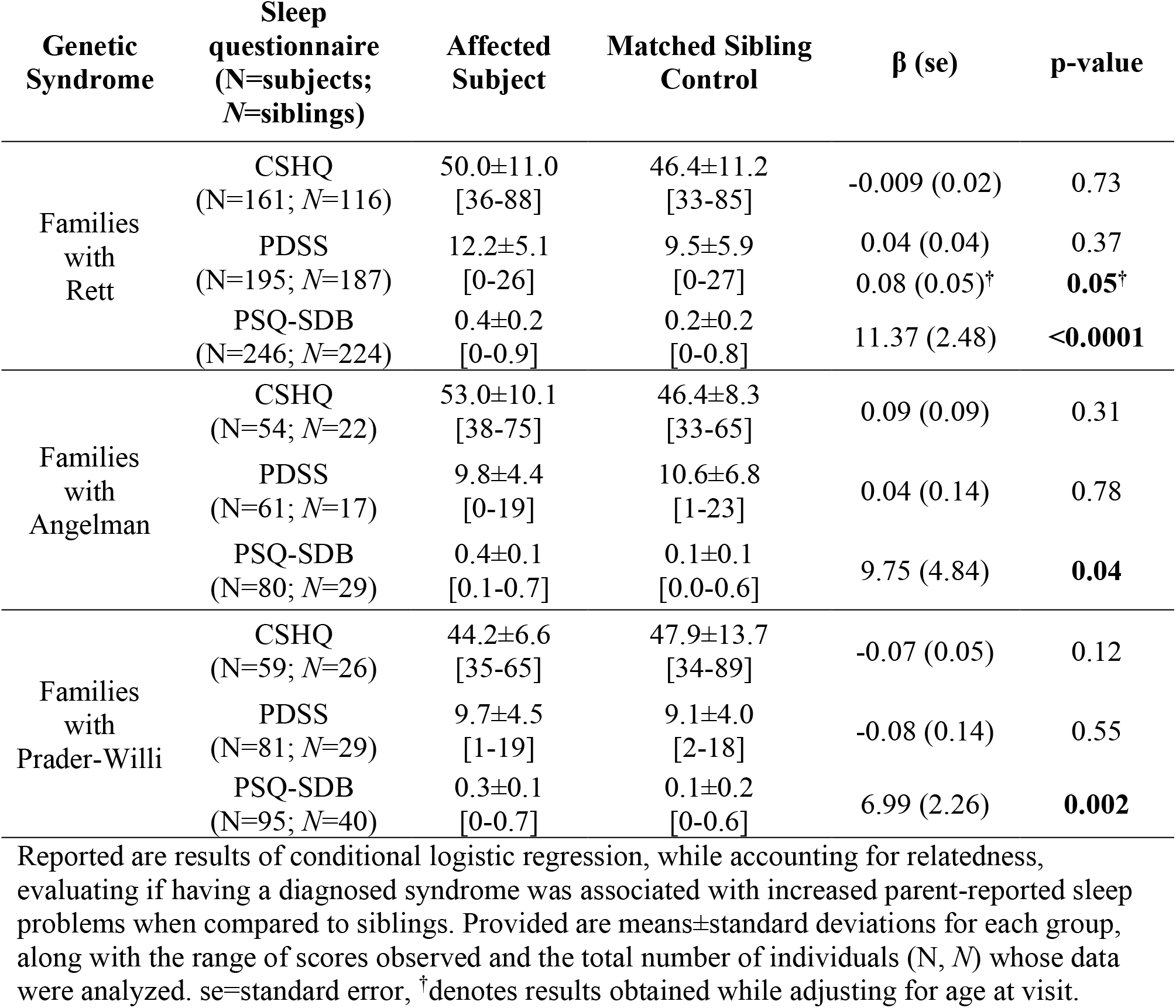
Severity of sleep problems in genetic syndromes compared to sibling controls.

## Conclusions

This study focused on characterizing patterns of sleep abnormalities in children and adolescents with RTT, AS, and PWS, including a comparison with their typically developing siblings. Participants were recruited from the RDCRN and were comprehensively characterized in terms of molecular diagnosis and clinical characteristics. By leveraging the resources of the RDCRN, we were able to analyze sleep data from one of the largest cohorts of individuals affected with these rare neurodevelopmental syndromes recruited to date. Our results suggest that sleep problems occur frequently in these syndromes; they also recapitulated evidence from previous studies conducted in smaller datasets but including objective sleep measurements.

Problems with sleep-disordered breathing were considered by parents to be more severe and problematic in individuals with these syndromes than in their typically developing siblings. Notably, individuals with RTT were reported to have more evidence of sleep-disordered breathing when compared to individuals with PWS. It is possible this reflects central sleep apnea in RTT, while obstructive sleep apnea is likely more prevalent in PWS given the relationship between PWS and obesity^31,41^, and the fact that obesity is a strong risk factor for obstructive sleep apnea^42^. In general, previous studies characterizing sleep patterns in RTT which have utilized objective measures, such as overnight polysomnography and wrist actigraphy, observed difficulties initiating and maintaining nocturnal sleep and early morning awakenings and not reported sleep-disordered breathing^43–45^. It is important to note that these studies were based on small sample sizes, between 9 and 20 participants with RTT, which could limit the scope of sleep disturbances observed and may not reflect those present in the majority of patients with RTT. Furthermore, respiratory disturbances are well-documented in RTT, although many of these disturbances (e.g., alternating bouts of breath-holding and irregular hyperventilation) are thought to be more pronounced during wakefulness; supporting a central cause^46^. Evidence also indicates that although breathing may be more irregular during the day, children with RTT demonstrate nocturnal breathing abnormalities^47^. Understanding sleep-disordered breathing in rare genetic syndromes is an important area for future work.

Our results indicate additional differences in the occurrence and severity of sleep problems among these genetic syndromes. For example, young children with AS have significantly more severe overall sleep problems compared to young children with RTT or PWS based on the total scores from the CSHQ. This finding is in agreement with previous studies suggesting sleep problems in individuals with AS are most severe in early childhood^48^. It is notable that we did not analyze subscale item scores reflecting the types of sleep problems in the CSHQ. Using CSHQ subscale scores to better define the specific types of problems that are unique to younger children with each syndrome is an area of interest to future studies. Middle-school age children and adolescents with RTT were reported to have more issues with daytime sleepiness on the PDSS compared to similarly aged individuals with AS or PWS. It is possible that this indicates issues with daytime sleepiness are influenced by female gender as RTT primarily affects females^49^. It is also possible that sleep-disordered breathing in patients with RTT influences expression of daytime sleepiness. However, of all the sleep traits evaluated, only the association between age and daytime sleepiness was significant and sleep-disordered breathing was not associated with age suggesting these two sleep-related issues may be distinct. In is also notable that the association between daytime sleepiness and age was unique to families with RTT. A previous study based on parent observations found the amount of total 24-hour sleep did not correlate with age, while the amount of daytime sleep positively correlated with age and the amount of night-time sleep negatively correlated with age^44^. In contrast, a study based on 24-hour wrist actigraphy found no correlation between the amount of sleep and age^45^. Our study provides additional support for an association between daytime sleep and age in RTT. Furthermore, the association between daytime sleepiness and a diagnosis of RTT (compared to their siblings) was near significance (p=0.05) when adjusting for age. These findings add to, and substantiate, a previous report which observed that individuals with RTT appear to sleep more during the day and to have a greater total sleep time in a 24-hour period compared to typically developing individuals^50^.

Sleep problems were prevalent in siblings of children with RTT, AS and PWS. This suggests that typically developing children living in households with children having these three developmental disorders may also experience more sleep problems than other children. Evidence from the CSHQ indicated that 81% of children between 2-10 years old who were diagnosed with RTT, AS or PWS had a total score ≥41 which is considered significant evidence of sleep problems. The prevalence of total CSHQ scores that were ≥41 in sibling controls was 67%. The prevalence of total sleep problems for both children with syndromes and their siblings was substantially greater than the currently reported prevalence of 20-40% for typically developing children^51,52^. This indicates that sleep behaviors in children with neurodevelopmental syndromes substantially influence those of their siblings, and perhaps also other household members, which may negatively impact the quality of life of both affected children and their families. These findings emphasize the importance of screening for sleep problems in the siblings of individuals with rare neurogenetic disorders as well as the individuals themselves.

Limitations of this study include lack of correlation of the findings from questionnaires with data from objective testing. Limited information regarding objective evaluations utilizing overnight sleep studies and actigraphy has been reported^50,53^. The use of objective assessments in future research may further characterize and quantitate sleep-disordered breathing and sleep/wake patterns in these patients. In addition, review of current or past sleep medications and information from the families concerning sleep medications was not obtained. While some medication data are available^54,55^, these data are limited and beyond the scope of the current study. Comprehensive characterization of the influences of various medications on expression of sleep problems is an area of interest for subsequent work. Furthermore, as the purpose of this initial study was to examine sleep problems, the relationship of comorbidities impacting sleep (e.g., epilepsy, gastrointestinal problems, clinical severity) was not evaluated and is also of interest for future work. Regardless of these shortcomings, our study provides valuable information about RTT, AS and PWS, and typically developing siblings that may contribute to clinical management of affected individuals and to the design of future studies evaluating the effectiveness of behavioral and pharmacological treatment of sleep problems, including clinical trials of novel interventions for these disorders.

## Funding

Supported by NIH U54 grants RR019478 (NCRR) and HD061222 (NICHD), IDDRC grants HD38985 (NICHD) and HD08321 (NICHD) and grant LM012870 (NLM). The Angelman, Rett and Prader-Willi Consortium (U54 RR019478 (NCRR) and HD061222 (NICHD)) is a part of the National Institutes of Health (NIH) Rare Disease Clinical Research Network (RDCRN), supported through collaboration between the NIH Office of Rare Diseases Research (ORDR) at the National Center for Advancing Translational Science (NCATS), and the Eunice Kennedy Shriver National Institute of Child Health and Human Development (NICHD). The content is solely the responsibility of the authors and does not necessarily represent the official views of the National Institutes of Health.

## Informed Consent Statement

All study participants or their legal guardian provided informed written consent prior to study enrollment.

## Author Disclosures

Alan K. Percy: PI of the NICHD-funded Natural History Study. Dr. Percy is site PI of clinical trials in Rett syndrome sponsored by Anavex and Acadia Pharmaceuticals and the investigator-initiated trial using ketamine. He is a consultant with Acadia. W.E.K. is Chief Medical Officer of Anavex Life Sciences Corp. He has also been consultant to AveXis, EryDel, GW, Marinus, Neuren, Newron, Ovid, Stalicla, and Zynerba, for which he has no relevant financial interest. L.M.B. has served as a paid consultant to pharmaceutical companies regarding clinical trial design for Angelman syndrome, as well as received money from pharmaceutical companies to conduct clinical trials in Angelman syndrome. None of this work involved sleep.

### Institutions where data analyses were performed

Baylor College of Medicine, Houston, TX; University of Alabama at Birmingham; University of South Florida; University of Kansas Medical Center, Kansas City, KS, USA

